# Negative Urgency as a Risk Factor for Hazardous Alcohol Use: Dual Influences of Cognitive Control and Reinforcement Processing

**DOI:** 10.1101/2021.05.02.442343

**Authors:** Eric Rawls, Noah R. Wolkowicz, Lindsay S. Ham, Connie Lamm

**Author notes:** Authors contributed equally and share joint first authorship (Eric Rawls, Noah R. Wolkowicz). Correspondence concerning this article should be addressed to: Eric Rawls, 717 Delaware St. SE, Minneapolis, MN 55414., Noah R. Wolkowicz, 216 Memorial Hall, Fayetteville, AR 72701.

## Abstract

Negative Urgency (NU) is a prominent risk factor for hazardous alcohol use. While research has helped elucidate how NU relates to neurobiological functioning with respect to alcohol use, no known work has contextualized such functioning within existing neurobiological theories in addiction. Therefore, we elucidated mechanisms contributing to the NU–hazardous alcohol use relationship by combining NU theories with neurobiological dual models of addiction, which posit addiction is related to cognitive control and reinforcement processing. Fifty-five undergraduates self-reported NU and hazardous alcohol use. We recorded EEG while participants performed a reinforced flanker task. We measured cognitive control using N2 activation time-locked to the incongruent flanker stimulus, and we measured reinforcement processing using the feedback-related negativity (FRN) time-locked to better-than-expected negative reinforcement feedback. We modeled hazardous drinking using hierarchical regression, with NU, N2, and FRN plus their interactions as predictors. The regression model significantly predicted hazardous alcohol use, and the three-way interaction (NU×N2×FRN) significantly improved model fit. In the context of inefficient processing (i.e., larger N2s and FRNs), NU demonstrated a strong relationship with hazardous alcohol use. In the context of efficient processing (i.e., smaller N2s and FRNs), NU was unrelated to hazardous alcohol use. This analysis provides preliminary evidence that brain mechanisms of cognitive control and reinforcement processing influence the relationship between NU and hazardous alcohol use, and confirms a specific influence of negative reinforcement processing. Future clinical research could leverage these neurobiological moderators for substance misuse treatment.

## 1 Introduction

Hazardous alcohol use is an ongoing public health issue, with a concerning proportion of individuals misusing or suffering from an alcohol use disorder (SAMSHA, 2018; WHO, 2018). One prominent risk factor for alcohol misuse is Negative Urgency (NU), the tendency to act rashly during negative emotional states (Berg et al., 2015; G. T. Smith & Cyders, 2016). Emerging work suggests NU is influential in neural responding related to impulse control (Chester et al., 2016; Gunn & Smith, 2010) and to affectively-laden reinforcers (Albein-Urios et al., 2013; Billieux et al., 2010; Chester et al., 2016; Cyders et al., 2015). This corresponds to research illustrating robust, positive associations between NU and alcohol use variables such as craving and use (VanderVeen et al., 2016). While research has helped elucidate how NU relates to neurobiological functioning with respect to alcohol use [(Chester et al., 2016; Cyders et al., 2014), reviewed in (Um et al., 2019)], no known work has contextualized such functioning within existing neurobiological theories in addiction (Goldstein & Volkow, 2002, 2011; Zilverstand et al., 2018). Understanding the NU – alcohol misuse relationship in the context of neuroscience addiction theories could lend insight into how NU promotes alcohol misuse, and provide precise intervention targets.

### 1.1 NU’s Behavioral and Neurobiological Correlates and its Relationship to Hazardous Alcohol Use

NU was delineated from a factor analysis of the Five Factor Model, yielding four psychological processes that lead to impulsive behaviors: Urgency, (lack of) Perseverance, (lack of) Premeditation, and Sensation Seeking (Whiteside & Lynam, 2001). Urgency was later split into Positive and Negative Urgency (Cyders & Smith, 2008b), referring respectively to tendencies for rash action during positive and negative moods. NU especially associates positively with general and heavy alcohol use, as well as alcohol-related consequences (Anthenien et al., 2017; Berg et al., 2015; Grimaldi et al., 2014; G. T. Smith & Cyders, 2016). Additionally, NU appears to demonstrate its effects only during negative emotional states or aversive outcomes (Cyders & Smith, 2007, 2008a, 2008b; VanderVeen et al., 2016), which emphasizes the role of negative reinforcement in understanding NU’s risk process.

Investigations into *how* NU imparts risk implicate cognitive control and stimulus appraisal. For example, NU is associated with poorer executive working memory capacity and the two interact to predict inhibitory control deficits (Gunn & Smith, 2010). NU is also associated with poorer inhibition of prepotent responding in the context of emotionally-valenced cues and greater risky decision-making in gambling paradigms (Billieux et al., 2010). NU is associated with reduced orbitofrontal cortex (OFC) activation in response to emotionally valenced stimuli (Cyders et al., 2015), and increased amygdala activation during the reappraisal and maintenance of negative emotions (Albein-Urios et al., 2013). Additionally, NU is associated with inefficient use of regions important for cognitive control (i.e., lateral prefrontal cortex) and habit-based behavior (i.e., dorsal striatum) during exposure to negatively valenced stimuli (Chester et al., 2016). This suggests that NU broadly predicts increases in neural responses to the need for cognitive control, and orients stimulus appraisal towards emotionally-relevant stimuli. Cognitive control and stimulus appraisal thus appear important in determining how NU imparts risk.

The pattern of relationships between NU, cognitive control, and stimulus appraisal are exemplified when considering NU’s associations with alcohol misuse. For example, higher levels of NU associate with greater alcohol craving in negative emotional states, greater alcohol seeking, and greater amounts of alcohol self-administration (VanderVeen et al., 2016). NU also predicts robust increases in expectations for positive alcohol outcomes (Fischer et al., 2012; Gunn & Smith, 2010; Spillane & Smith, 2010) and motivations to use alcohol as a means of alleviating negative moods and/or enhancing current moods (Adams et al., 2012; Curcio & George, 2011; Watkins et al., 2015). During exposure to alcohol cues, greater levels of NU predict greater activation in neural regions associated with attributing reward value, emotion-based decision-making (Cyders et al., 2014), and allocating and directing attentional resources (Chester et al., 2016). Further, insula activation for individuals with higher levels of NU predicts greater alcohol misuse at 1-month and 1-year follow ups (Chester et al., 2016); this is notable, given the insula’s implicated role in salience processing (Menon & Uddin, 2010).

Collectively, such research suggests *how* an individual processes and responds to affectively-laden reinforcers, such as alcohol for misusers, may play a role in NU’s risk impartation process. That is, individual differences in the efficiency and extent to which negative reinforcement is processed may significantly increase the expression of NU’s rash action. This reasoning aligns with contemporary neuroscientific models on the underpinnings of addiction, which emphasize the role of cognitive control and reinforcement in the initiation and maintenance of addiction.

### 1.2 Neurobiological Dual Models of Addiction

Dual models of drug addiction (Goldstein & Volkow, 2002, 2011; Zilverstand et al., 2018), focus on the role of reinforcement and cognitive control processing in the cycle of addiction. Following initial substance use, activation patterns change in frontal cortical structures (responsible for cognitive control, including conflict monitoring and inhibition of inappropriate responses) and limbic structures (responsible for incentivizing substance-related reinforcers and disincentivizing alternative reinforcers). Reductions in cognitive control can be exacerbated by changes in reinforcement salience processing, pointing to a likely interaction between processes of cognitive control and reinforcement processing. In dual models of addiction, cognitive control is broadly construed and is operationalized using flanker and Stroop tasks (which require withholding an inappropriate response and selecting an alternate response) and go/no-go and stop signal tasks (which require withholding an inappropriate response without selection of an alternate response) (Goldstein & Volkow, 2011; Zilverstand et al., 2018). Salience or reinforcement processing describes the increased valuation of drug rewards and the devaluation of non-drug rewards in addiction; increased valuation of drug rewards is generally assessed using cue exposure paradigms while decreased valuation of non-drug rewards is typically assessed using paradigms that deliver point feedback as a proxy for monetary rewards, such as the Monetary Incentive Delay task (Knutson et al., 2001).

In measuring the two components of dual models of addiction, prefrontal cortex (PFC) activity provides a reliable marker of neural alterations in substance abuse (Goldstein & Volkow, 2011). PFC is a primary target of midbrain dopaminergic projections (Floresco & Magyar, 2006; Lammel et al., 2011; Smith et al., 2014), and is critical for the integration of cognitive control and reinforcement (Lammel et al., 2014; Xu and Yao, 2010). Dual processes of incentive salience processing and cognitive control are important for decision making (Cools, 2016; Yee & Braver, 2018), and dynamically contribute to an individual’s decision to engage in or abstain from impulsive behavior (Gladwin et al., 2011; Luna et al., 2013; Westbrook & Frank, 2018). PFC activity is therefore a critical hub underlying maladaptive cognitive control and reinforcement processing in substance abuse.

ERPs (averaged EEG) linked with PFC function can directly measure cortical activity underlying cognitive control and salience processing. ERPs have utility as markers of individual differences in cortical efficiency (Denke et al., 2018; Jabr et al., 2018; Lamm et al., 2014, 2018; Linka et al., 2005; McDermott et al., 2009; Meyer et al., 2014, 2017; Rawls et al., 2018; Troller-Renfree et al., 2018). The N2 is a mediofrontal ERP component tied to cognitive control processes (Bekker et al., 2005; Jonkman, 2006; van Veen & Carter, 2002). N2 activation marks impaired cognitive control in substance abuse (Buzzell et al., 2014; Luijten et al., 2011), and has been used to link trait-level predispositions to outcomes associated with lack of cognitive control (Denke et al., 2018; Lamm et al., 2018; Rawls et al., 2018). Note that while P3 activation has also been linked to resolution of cognitive control requirements (Wessel & Aron, 2015), we have previously shown that P3 activation predicts congruent trial responses times more strongly than incongruent response times (Rawls, Miskovic, & Lamm, 2020). As such, the N2 likely provides a more specific metric for quantifying conflict monitoring in flanker-type tasks (such as that employed in the current study). The feedback-related negativity (FRN; alternately called the reward positivity [RewP]) is a related mediofrontal potential that occurs following reinforcement feedback. The FRN/RewP reflects prediction errors (Holroyd & Coles, 2002), a common currency underlying reinforcement (Averbeck & Costa, 2017; Gläscher et al., 2010; Glimcher, 2011). Recent evidence has demonstrated that the FRN/RewP encodes continuous PE magnitudes for reinforcing outcomes in both positive and negative reinforcement conditions, but with opposite sign (Hird et al., 2018; Rawls, Miskovic, Moody, et al., 2020; Rawls & Lamm, 2021; Sambrook & Goslin, 2014; Soder & Potts, 2018; Talmi et al., 2013).

### 1.3 Current Study

We examined the influence of neural measures of impaired cognitive control and reinforcement processing, as potential mechanisms tying NU to hazardous alcohol use. As it relates to NU, the emphasis of dual models of addiction on cognitive control and reinforcement processing highlight underlying neural processes that during negative reinforcement, may increase tendencies for emotionally-dependent disinhibition and substance-related stimuli preference. Individuals with trait-level tendencies prioritizing the alleviation of, and who demonstrate poor top-down control when experiencing, aversive stimuli may be particularly vulnerable for engaging in risky behaviors (including hazardous alcohol use) if their neural functioning also exacerbates these tendencies. Unfortunately, no known work has assessed how neural mechanisms of cognitive control and reinforcement processing impact NU’s connection to such risky behaviors. This is an important gap in the literature, given the prominence of dual models in neuroscientific accounts of addiction, and the prominence of NU in empirical accounts of hazardous alcohol use. We hypothesized that the relationship between NU and hazardous alcohol use would be stronger for individuals who 1) demonstrated inefficient neural processing of cognitive control (i.e., inefficient/more negative N2 activation), 2) demonstrated inefficient neural processing of reinforcement (i.e., inefficient/more negative FRN activation), and 3) the relationship between NU and hazardous alcohol use would be strongest for those who demonstrate inefficient control *and* reinforcement processing, while this relationship would be weakest for those who demonstrate both efficient cognitive control and reinforcement processing.

## 2 Methods and Materials

### 2.1 Participants

Ethical approval for this study was obtained from the University of Arkansas IRB (#1708016049). Fifty-five undergraduates (37 female, *M_age_* = 19.2±2.06, 2 left handed) completed the study after giving informed consent. Participants were excluded based on self-reported psychiatric diagnoses or uncorrected visual impairments. These data were previously reported in Rawls and Lamm (2021). In that study we report single-trial sensitivity of FRN/RewP amplitude to the magnitude of negative reinforcement prediction errors (an aversion positivity). Flanker-locked ERPs, NU, and AUDIT data reported in this study have not been previously reported. See *Table 1* for full demographics and descriptive statistics on the participant group.

**Table 1.** Demographics and descriptive statistics for the current sample.

| Variable | Descriptives |
| --- | --- |
| **Gender** | 37 female, 18 male |
| **Race** | 2 Asian, 1 Black, 42 White, 4 Other, 6 Multiple |
| **Ethnicity** | 6 Hispanic/LatinX, 49 not Hispanic/LatinX |
| **Age** | Mean = 19.2, SD = 2.06, Min = 18, Max = 30 |
| **Negative Urgency** | Mean = 2.08, SD = 0.76, Min = 1, Max = 3.83 |
| **Response Inhibition (N2)** | Mean = 0.61, SD = 2.94, Min = -6.34, Max = 7.62 |
| **Reinforcement processing (FRN)** | Mean = 0.21, SD = 1.36, Min = -3.18, Max = 4.98 |
| **AUDIT sum** | Mean = 13.09, SD = 5.14, Min = 1, Max = 25 |

### 2.2 Measures and Materials

#### 2.2.1 UPPS Impulsive Behavior Scale

NU was assessed using the UPPS-P Impulsive Behavior Scale (Lynam et al., 2006; Whiteside & Lynam, 2001). The UPPS-P is a 4-point Likert scale (1 = *agree strongly*, 4 = *disagree strongly*) self-report measure that assesses the five components of the UPPS-P impulsivity model. This measure has a well-validated factor-structure (Cyders et al., 2007) and the NU scale showed excellent reliability in the present sample, α = .94.

#### 2.2.2 Alcohol Use Disorders Identification Test (AUDIT)

The AUDIT is a self-report questionnaire assessing past-year alcohol use, dependence symptoms, and related problems (Kokotailo et al., 2004). Items are rated on 5-point or 3-point scales and summed to produce a total score. The measure is well-validated for use with collegiate populations (Kokotailo et al., 2004) and demonstrated high reliability in the present sample, α = .85. We did not have any inclusion or exclusion criteria for participant drinking behaviors, as our purpose was to examine determinants of drinking behavior in a normative collegiate sample. Importantly, our sample demonstrated normative drinking behaviors on the AUDIT scale, M = 13.09 ± 5.14 (minimum = 1, maximum = 25), and all participants demonstrated at least some level of alcohol use (AUDIT ≥ 1).

#### 2.2.3. Reinforced flanker task

Participants completed a reinforced flanker task, presented using E-Prime 3.0 (Psychology Software Tools, Pittsburgh, PA), and initially reported in Rawls and Lamm (2021) (*Figure 1*). This task provided a means of assessing neural responding with respect to cognitive control and reinforcement processing and was thus ideal for our study’s aims of assessing how these components of dual models of addiction (i.e., cognitive control and reinforcement processing) influenced the relationship between NU and hazardous alcohol use.

**Figure 1.**
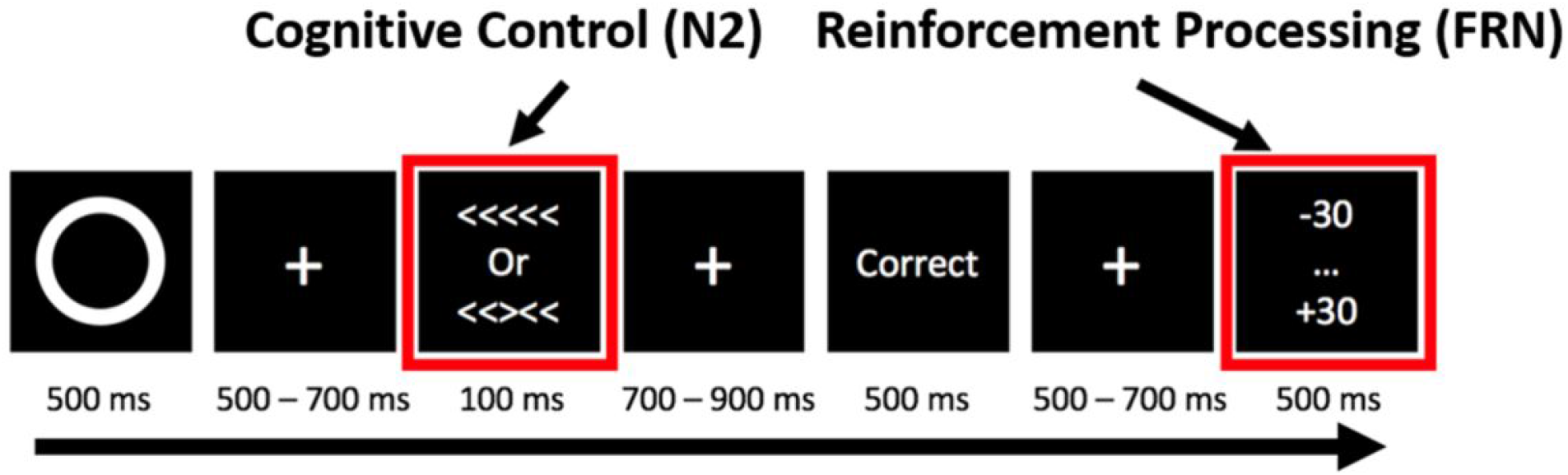
Task diagram for the modified reinforcement flanker paradigm. Participants were cued as to whether the current trial was to be positive or negative reinforcement with a white square or a white circle, respectively. Negative reinforcement trials are of primary importance for the current manuscript. Participants then had to respond to a congruent or incongruent flanker arrow stimulus. Only correct trials were analyzed, and participants were shown the correct feedback image on every trial prior to the point feedback being shown. Finally, participants were given some amount of points (on average +0 for negative reinforcement). The crucial task manipulation was that for every trial, the amount of points given deviated slightly from the overall mean expectation, generating prediction errors in better-than-expecteds/worse-than-expected outcomes. ERP values were separately computed for better-than-expected outcomes (+11 to +30 points above expectation) and worse-than-expected outcomes (−30 to −11 points below expectation); outcomes that were roughly as-expected (−10 to +10 points from expectation) were not analyzed.

The trial structure of the task was as follows: first, a fixation was presented before every trial. A black-and-white square or circle, denoting positive or negative reinforcement trials respectively (Knutson et al., 2001), was presented for 500 ms, followed by a fixation that lasted 500 – 700 ms. In the current analysis, we focus our attention on the negative reinforcement condition, given the link between NU and negative reinforcement processes. Congruent (< < < < < or > > > > >) or incongruent (< < > < < or > > < > >) arrows were shown for 100 ms, followed by a fixation lasting 900 – 1100 ms. Correct/incorrect feedback was shown for 500 ms (only correct trials were analyzed, to avoid influences of error monitoring), followed by a fixation lasting 500 – 700 ms. Point feedback was shown for 1000 ms. Average return for correct negative reinforcement trials was zero points, but actual return varied from −30 to +30 points. Before participants began, they completed 50 trials for practice. During practice trials, the outcome was always as expected (correct answers resulted in no loss or gain of points). Participants progressed to the main task after obtaining 80% or better accuracy during practice. The task contained 960 trials divided into 16 blocks of 60 trials, requiring approximately 90 minutes.

In line with previous research using random (as opposed to blocked) reinforcement presentations (Knutson et al., 2001), participants were forewarned about the designation of trials as either positive or negative reinforcement. This created an expectation of the potential outcomes of wins or losses for each trial. By creating an expectation of average outcome in the practice and then systematically providing more or less points than expected on correct trials, this task produced better-than-expected and worse-than-expected outcomes without the confounding effect of error monitoring. Therefore, while all trials were “wins” in the sense that “correct” feedback was delivered, trials could be better-than-expected or worse-than-expected. While we recorded both EEG and behavior during task performance, the current analyses only concerned the EEG recording, as behavior only occurred in response to the flanker stimulus. Therefore, behavioral indices captured only processing of cognitive control, while EEG captured processing of both cognitive control and reinforcement processing.

### 2.3 EEG processing

EEG data were acquired and preprocessed as described in Rawls & Lamm (2021). EEG were sampled at 1000 Hz using a 129-channel EGI net referenced to vertex (Philips EGI, Inc.). Recording began after impedances were reduced below 50 kΩ. Data were processed using EEGLAB 15 (Delorme & Makeig, 2004) and MATLAB 2017 (Mathworks). Data were downsampled to 125 Hz and filtered between 0.1 Hz and 30 Hz using zero-phase FIR filters. Channels were removed based on SD >= 4. Copies of datasets were high-pass filtered at 1 Hz (Winkler et al., 2015) and epoched into 3 second windows surrounding point feedback onset or flanker onset (from −1 s before to 2 s after). Infomax ICA (Makeig et al., 1996) was computed on the 1 Hz filtered dataset. Likely artifactual independent components were detected using the SASICA plugin (Chaumon et al., 2015) using autocorrelation statistics, focal component activity, and the ADJUST plugin (Mognon et al., 2011). ICs were additionally labeled using ICLabel (Pion-Tonachini et al., 2019). ICs labeled as eye, heart, muscle, line noise, or channel noise with greater than 70% confidence were marked as artifactual. ICA weights and artifact components calculated in the 1 Hz high-pass filtered dataset were copied to the 0.1 Hz high-pass filtered dataset, and detected artifactual ICs were visually verified and removed from the data; all further analyses were performed on the 0.1 Hz filtered dataset.

EEG were epoched time-locked to the flanker stimulus and the point feedback. Epochs were baseline corrected for 200 ms prior to stimulus presentation and threshholded at ±125 μV, removed channels were interpolated using spherical splines, and data were re-referenced to the montage average. N2 amplitude was calculated as the mean voltage from 250 – 350 ms post-stimulus (flanker) at a mean of three mediofrontal sensors (Fz, FCz, Cz), and FRN amplitude was calculated as the mean voltage from 250 – 350 ms post-feedback (point outcome) at a mean of three mediofrontal sensors (Fz, FCz, Cz). These scalp locations and time windows were selected based on prior literature, and we inspected grand-average waveforms over the scalp to confirm that these scalp locations and time windows contained the relative negativities associated with the N2 and FRN.

Given NU’s connection to rash action potentially in relief of aversive stimuli, only negative reinforcement trials were considered for the current analysis. For the FRN, since outcomes varied continuously, we split trials into three conditions for analysis as individual difference variables. Actual outcomes varied from −30 to +30 points, with an average (expected) return of 0 points. We took trials with outcomes ranging from −30 to −11 points as worse-than-expected trials, and trials with outcomes ranging from +11 to +30 points as better-than-expected trials (trials with outcomes ranging from −10 to +10 points were considered expected outcomes [not analyzed]). The average number of trials making up each ERP (Mean ± SD) were as follows: negative reinforcement better-than-expected FRN = 119.54 ± 21.37, negative reinforcement incongruent N2 = 150.38 ± 43.05. ERP waveforms and scalp topographies are depicted in *Figure 2*.

**Figure 2.**
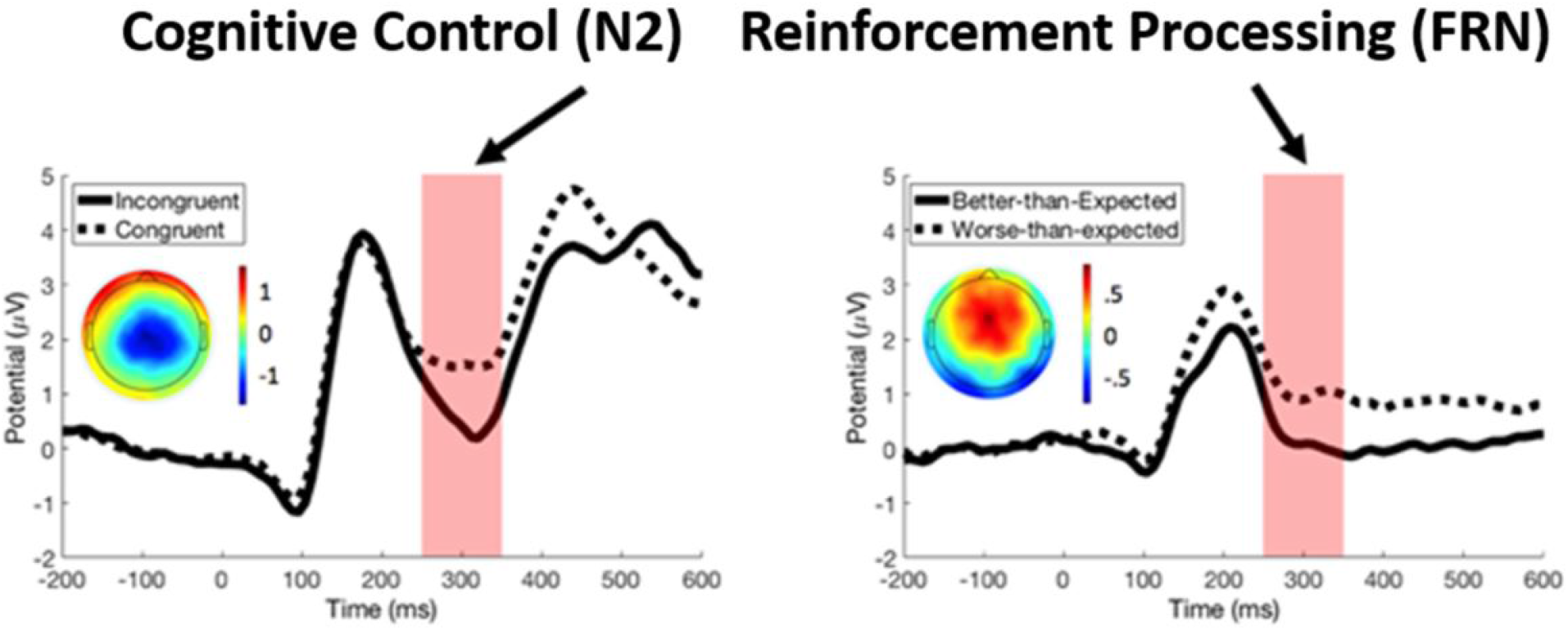
Grand-average ERP waveforms derived from the modified reinforcement flanker task. Note that positive values are plotted upward on the y-axis. Left panel indicates the ERP time-locked to the flanker arrow stimulus and separated by stimulus congruency, and the topographic plot illustrates the scalp distribution of the incongruent – congruent difference waveform from 250 to 350 ms post-stimulus. Right panel indicates the ERP time-locked to negative reinforcement feedback presentation and separated into better-than-expected and worse-than-expected outcomes (outcomes that were roughly as expected were not analyzed), and the topographic plot illustrates the scalp distribution of the worse-than-expected – better-than-expected difference. The incongruent N2 was selected as a measure of cognitive control and the better-than-expected FRN was selected as a measure of reinforcement processing. Primary moderation analyses examined the raw condition ERPs as moderators, however, we also ran control analyses using residualized ERP amplitudes (incongruent N2 with congruent N2 regressed out, better-than-expected FRN with worse-than-expected FRN regressed out).

### 2.4 Predicting AUDIT scores as a function of NU and iRISA

To determine if impaired cognitive control and reinforcement processing in combination with NU provide a better predictor of hazardous alcohol use than NU alone, we used multiple regression implemented in the PROCESS v3.3 macro (Hayes, 2017) and SPSS 26. We considered ERP measures of control and reinforcement as moderators (Denke et al., 2018; Jabr et al., 2018; Lamm et al., 2014, 2018; Meyer et al., 2017; Rawls et al., 2018) of the relationship between NU and AUDIT scores. For a diagram of the proposed model see *Figure 3*. We use moderation analysis to test the hypothesis that the relationship between NU and hazardous alcohol use depends on neural mechanisms underlying cognitive control and reinforcement processing. Dual models of addiction suggest that cognitive control and reinforcement processing interact. Specifically, cognitive control of inappropriate actions weakens when reinforcement salience is high (Goldstein & Volkow, 2011). This interactive effect is best captured by a moderation model. Therefore, a moderated-moderation model was examined using PROCESS model 3. Based on the theoretical link between NU and rash action specifically undertaken in attempts to relieve aversive states, we examined neural markers of cognitive control and reinforcement processing from negative reinforcement trials as moderators in this analysis. However, to test whether our results are specific to negative reinforcement contexts, we also examined ERPs from positive reinforcement trials. Variables were standardized prior to regression analysis (Dawson, 2014).

**Figure 3.**
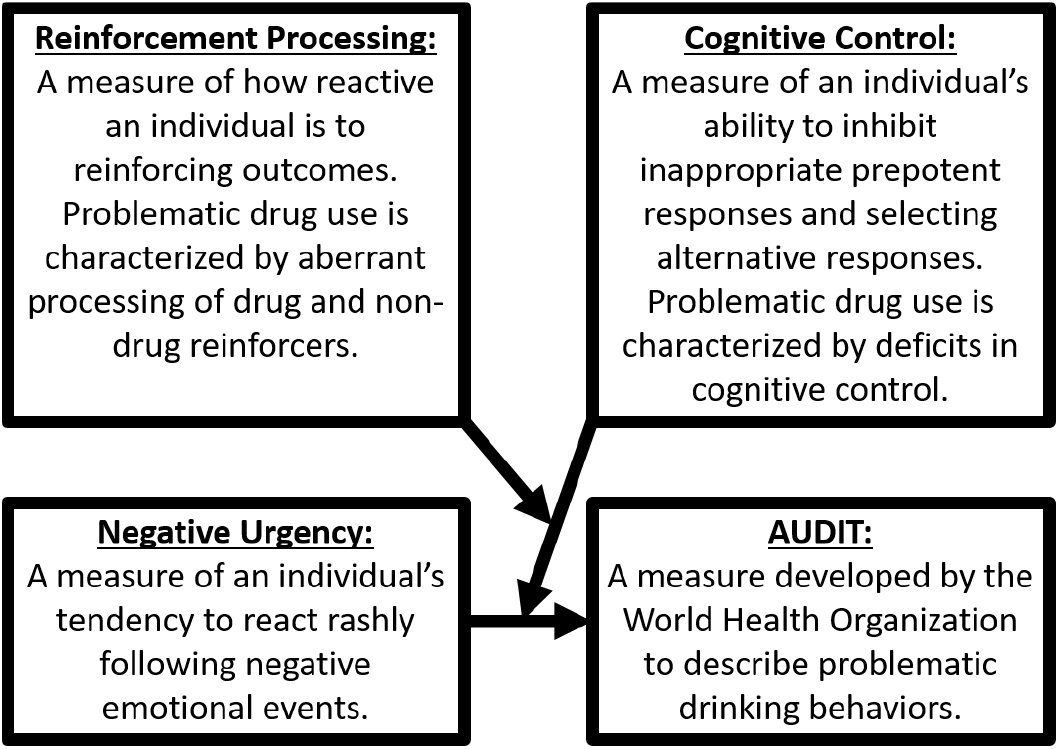
Model diagram for the proposed moderated-moderation model. We suggest that neural indices of both impaired cognitive control and impaired reinforcement processing should moderate the relationship between individual differences in NU and problematic drinking behaviors. This effect is in line with dual models of addiction, which do not suggest that cognitive control causes reinforcement processing, or that reinforcement processing causes cognitive control (the assumption of causation is necessary for a mediation analysis). Instead, dual models suggest that the influence of cognitive control on eventual actions can be weakened or eliminated entirely by overly inefficient reinforcement processing. The appropriate test, when the relationship between variables changes as a function of another variable, is a test for moderation.

Since power estimates and recommended sample sizes for moderated-moderation models are not well established, we computed bootstrapped confidence intervals for all parameters in the regression analysis (10,000 randomizations), allowing greater confidence in this analysis. We report both unadjusted *R^2^* and adjusted *R^2^*, providing an estimate of the variance explained that is adjusted for bias due to sample size and number of model parameters. We do not posit that NU causes brain activity underlying cognitive control and reinforcement processing, a necessary assumption of mediation (Hauser et al., 2014). Nonetheless, as some research indicates neurocognitive measures mediate (Chester et al., 2016; Cyders & Smith, 2008b) the relationship between NU and AUDIT scores, we tested whether our data met the assumptions for ERPs to function as mediators, rather than moderators as our model proposes.

This analysis used raw condition-specific ERP amplitudes as individual difference variables, to avoid the unfavorable properties of difference scores when analyzed as individual difference variables (Meyer et al., 2017). However, the recent report by Meyer and colleagues (2017) demonstrated that residualized ERP amplitudes might overcome some of the issues inherent in difference scores. Therefore, we also tested the reported regression model using the residualized N2 amplitude (incongruent N2 with congruent N2 regressed out) and the residualized FRN (better-than-expected FRN with worse-than-expected FRN regressed out) in place of the raw condition-specific scores. To examine whether our results were specific to negative reinforcement (as expected based on the theoretical link of NU to negative reinforcement processes) we also examined ERPs following positive reinforcement outcomes (S1).

## 3 Results

### 3.1 Preliminary Analysis: Potential Confounding Variables and Correlation Structure

Independent-samples *t*-tests were conducted on N2, FRN, NU, and AUDIT with gender as the independent variable. Results indicated no gender differences in any variables, all *p* > .39. Pearson correlations were computed between participant age and each variable of interest. Age was uncorrelated with any considered variable, all *p* > .22. Moderation analysis is appropriate when moderators are uncorrelated with the IV/DV, whereas mediation requires significant correlations between the mediator and IV (Baron & Kenny, 1986; Dawson, 2014). Correlation results (Table 2) indicated that N2 and FRN are uncorrelated with NU or AUDIT scores. NU was correlated with AUDIT sum.

**Table 2.** Preliminary correlation analysis revealed that neither proposed moderator was correlated with either NU or AUDIT scores, indicating that a moderation analysis is more suitable than a mediation analysis for examining the integration of dual models of addiction and NU theories. The two proposed moderator variables were highly correlated, which makes sense as they were brain potentials recorded from the same scalp location (albeit following different cognitive events). However, inspection of model tolerance values (VIF) indicated no problems with multicollinearity in any of the models (all VIF < 2.5) so this correlation is not a problem statistically. * = *p* < .05, ** = *p* < .01

|  | Negative Urgency (NU) | Cognitive control (N2) | Reinforcement processing (FRN) |
| --- | --- | --- | --- |
| **Negative Urgency (NU)** | - | - | - |
| **Cognitive control (N2)** | $r(54) = .05, p = .73$ | - | - |
| **Reinforcement processing (FRN)** | $r(54) = .02, p = .91$ | $r(54) = .42, p = .001^{**}$ | - |
| **AUDIT sum** | $r(54) = .30, p = .03^{*}$ | $r(54) = -.07, p = .60$ | $r(54) = .02, p = .90$ |

### 3.2 Neural Measures of Impaired Cognitive control and Reinforcement processing Moderate the Association Between NU and Problematic Drinking Behavior

A moderated-moderation model was run as hierarchical regression with NU, N2, and FRN entered as predictors on the first step, two-way interactions on the second step, and the three-way interaction (NU×N2×FRN) on the third step (model summary presented in *Table 3*). This analysis used ERPs derived from negative reinforcement trials, given the theoretical link between negative reinforcement processes and negative urgency. The model significantly predicted AUDIT scores, *F*(7,47) = 3.15, *p* = .008, unadjusted *R^2^* = .32 (adjusted *R^2^* = .22). The addition of a three-way interaction in the third step provided an additional unadjusted *R^2^* of .07 (adjusted *R^2^* of .07), indicating a significant increase in explanatory power, *F*(1,47) = 5.06, β = −2.19, 95% bootstrap CI = [−4.16, −.23], *t*(47) = −2.25, *p* = .03. Effects of NU on AUDIT scores were probed at −1 SD, +1 SD, and the mean value for both moderators (N2 and FRN). Moderating effects of impaired cognitive control (N2) were only significant when reinforcement processing (FRN) was high (+1 SD), representing efficient reinforcement processing. The positive relationship between NU and AUDIT score was attenuated when both cognitive control and reinforcement processing were efficient and exacerbated when they were inefficient. Simple slopes of probed moderating effects and significance tests for slopes are summarized in *Figure 4.*

**Figure 4.**
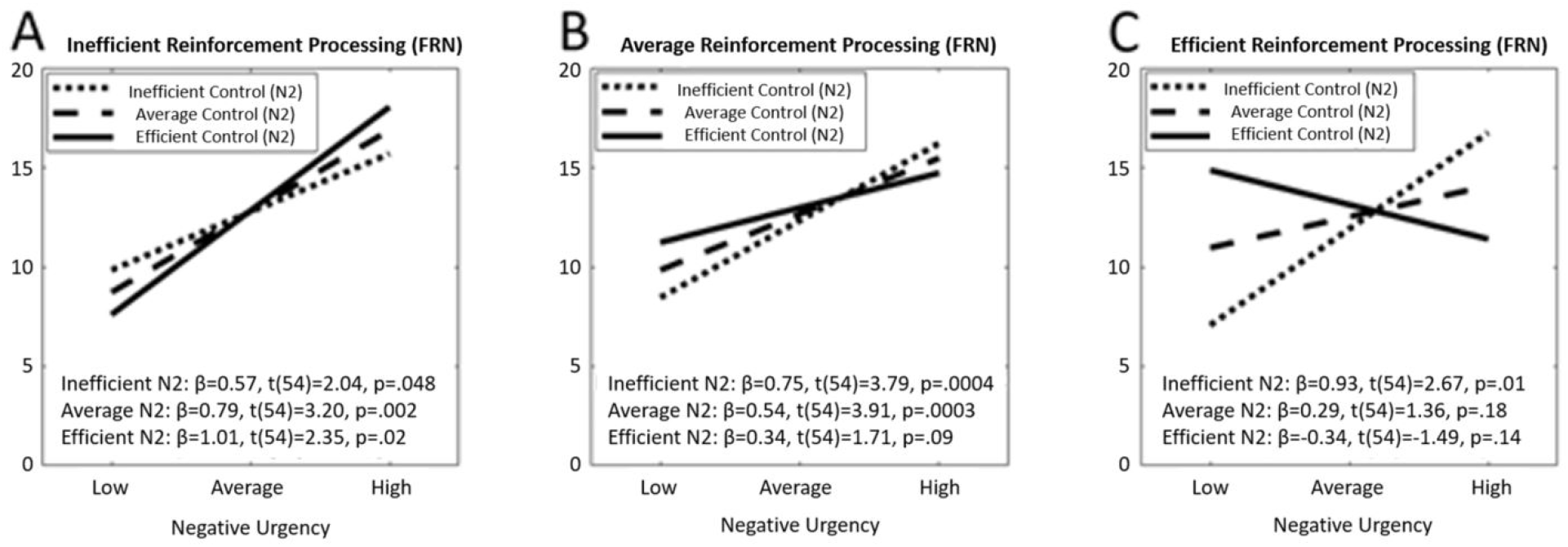
These moderated moderation results suggest a theoretical interpretation where control over prepotent actions moderates the influence of NU on hazardous alcohol use, but the moderating effect of prepotent cognitive control is itself moderated by reinforcement processing. This result shows that the moderating role of cognitive control is most present when paired with efficient reinforcement processing and is most clearly seen when comparing image “A” on the left to image “C” on the right. Moderation effects were probed at −1 SD, the mean, and +1 SD. Image “A” illustrates at inefficient levels of salience processing (−1 SD), a strong, positive relationship between NU and hazardous alcohol is present regardless of the efficiency for cognitive control. In contrast, image “C” illustrates when efficient levels of salience processing (+1 SD) are paired with efficient levels of cognitive control (+1 SD), NU’s relationship with hazardous alcohol use is almost reversed (and is non-significant). Thus, our analyses revealed that moderating effects of inhibition were most prominent when reinforcement processing was also efficient. Further, our analyses indicated the relationship between NU and AUDIT sum was weakest, and therefore psychiatric outcomes best, when both inhibition and reinforcement processing were efficient.

**Table 3.** To test the hypothesis that cognitive control and reinforcement processing moderate the impact of negative urgency on AUDIT scores, we employed a moderated-moderation regression analysis (PROCESS model 3) in SPSS. Results indicated a significant main effect of NU on drinking behavior, as well as the expected three-way interaction of NU × N2 × FRN.

| | Unstandardized $\beta$ | 95% CI | $t(55)$ | $p$-value |
| --- | --- | --- | --- | --- |
| **Negative Urgency (NU)** | 2.80 | [1.36, 4.23] | 3.91 | .0002*** |
| **Cognitive control (N2)** | 0.32 | [-1.27, 1.91] | 0.40 | .69 |
| **Reinforcement processing (FRN)** | -0.13 | [-1.66, 1.40] | -0.17 | .89 |
| **NU × N2** | -1.06 | [-2.51, .39] | -1.48 | .15 |
| **NU × FRN** | -1.28 | [-3.21, .64] | -1.34 | .19 |
| **N2 × FRN** | 0.29 | [-0.84, 1.42] | 0.52 | .61 |
| **NU × N2 × FRN** | -2.19 | [-4.16, -.23] | -2.25 | .03* |

To test the specificity of our results to negative reinforcement (as opposed to positive reinforcement), we also ran this moderated-moderation analysis using ERPs from positive reinforcement trials. In positive reinforcement, incongruent N2 moderated the relationship of NU with AUDIT scores, but FRN did not play any role in this relationship (S1). This important control analysis reveals the specific role of neural processing of negative reinforcement outcomes in shaping the relationship of NU to problematic drinking behaviors.

## 4 Discussion

### 4.1 General Discussion

Negative Urgency (NU) robustly predicts hazardous alcohol use. However, our understanding of how this trait construct initiates and maintains problematic drinking behaviors is still developing. Integrating NU’s influences on neurobiological functioning within existing neuroscientific theories, particularly dual models of addiction (Goldstein & Volkow, 2002, 2011; Zilverstand et al., 2018), is therefore essential, especially as research on structural and functional targets grows (Um et al., 2019). Our findings provide initial evidence that individual differences in cortical activation underlying cognitive control and reinforcement processing can either exacerbate or attenuate the association between NU and hazardous alcohol use. These results integrate theoretical perspectives on NU with neurobiological dual models of addiction and suggest future intervention targets.

### 4.2 Integrating Dual Models of Addiction into our Understanding of NU and Hazardous Alcohol Use

Dual models of addiction suggest that maladaptive cognitive control and reinforcement processing underlie drug seeking and taking behaviors (Goldstein & Volkow, 2002, 2011; Zilverstand et al., 2018). A recent update to this theory (Zilverstand et al., 2018) found impairments in salience network (including anterior insula and dorsal anterior cingulate cortex [ACC]) and executive network (including dorsolateral and ventrolateral PFC) throughout the entire addiction cycle, suggesting deficits in frontal cortical activation are present from an early time point in substance misuse. The N2 and FRN, evoked potentials peaking approximately 300 ms following conflict-related stimuli (N2) or reinforcing feedback (FRN), are related to neural generators in salience and executive networks. For example, an analysis of cognitive control in alcoholics used LORETA to localize N2 to anterior cingulate and prefrontal cortices (Pandey et al., 2012), and the FRN was modeled to generators in anterior cingulate cortex and other areas of the salience network (Hauser et al., 2014).

Our results indicate that prefrontal brain potentials reflecting cognitive control (N2) and reinforcement processing (FRN) exert interactive moderating effects that clarify the relationship of NU with maladaptive drinking behaviors. Specifically, brain activity suggesting efficient cognitive control moderates the relationship between emotion-based predisposition to rash action and levels of reported alcohol misuse. Simultaneously, neural activity indicating inefficient reinforcement processing moderates this moderation relationship, such that the relationship between NU and alcohol misuse is non-significant when both N2 and FRN activation are efficient. This result is in line with the iRISA model, which indicates that inefficient reinforcement processing can override cognitive control control (Goldstein & Volkow, 2011).

### 4.3 Clinical Insights and Potential Therapeutic Benefit

In line with the NIMH Research Domain Criteria (RDoC), these results provide preliminary evidence that neurobiologically-grounded theories can inform clinical practice and our understanding of clinically significant outcomes, such as alcohol misuse. Critically, our results suggest a model incorporating neural measures corresponding to dual models of addiction can provide more than three times the explanatory power of alcohol misuse in comparison to a model using only NU (itself a prominent clinical predictor of alcohol misuse). Thus, the combination of clinical and neurobiologically-informed theory in our research provide greater insight into the mechanisms that contribute to maladaptive alcohol use behaviors.

Our results lend insight into how clinicians might intervene to reduce the impact of trait-level characteristics on alcohol misuse. NU is empirically established as a risk factor for alcohol misuse, but little work explains how clinicians might translate such knowledge into effective interventions. Developing impactful interventions is facilitated by delineating specific psychophysiological targets that impact relationships between trait-level risk factors and alcohol misuse. Our results suggest interventions that enhance efficiency in an individual’s use of cortical resources for cognitive control and reinforcement processing (i.e., identification of salient stimuli and shifting of attention toward or away from such stimuli) may significantly impact their engagement in hazardous alcohol use. Importantly, previous research by our group has shown that PFC function might have a downstream causal impact on severity of alcohol use disorder (Rawls et al., 2021), suggesting that interventions focused on PFC function could improve cognitive function in individuals with problematic alcohol use.

To date, several researchers have developed interventions that may be relevant to these examples, targeting basic behavioral or brain processes through training or the use of neurofeedback. For example, attentional bias modification trainings indicated promise in improving alcohol misusing patients’ ability to disengage from alcohol-related cues and achieve at least short-term reductions in drinking (Cox et al., 2014; Schoenmakers et al., 2010). Research also suggests fMRI neurofeedback can produce improvements in alcohol-related cue-reactivity (Kirsch et al., 2016), emotion-regulation (Zilverstand et al., 2015; Zilverstand, Parvaz, et al., 2017), and attentional control (Zilverstand, Sorger, et al., 2017). However, no known work has assessed how targeting the efficiency of cortical processing as it relates to cognitive control and reinforcement processing (e.g., N2 and FRN amplitudes), may improve clinical outcomes in high NU and/or alcohol misusing individuals.

### 4.4 Limitations and Future Directions

Our study is the first to apply the iRISA neurobiological theory of addiction to test potential mechanisms underlying the link between NU and hazardous alcohol use, and therefore has several limitations. First, our study used a relatively small sample size of 55 participants. This is a high-powered study among ERP studies, but a relatively small sample compared to some clinical studies. While we used a similar sample size to previous studies that considered ERPs as moderators of relationships between predispositions and outcomes (Denke et al., 2018; Rawls et al., 2018, 2018), our analysis contained two moderators while previous studies have used only a single moderator. To overcome these limitations, we used bootstrapping (10,000 randomizations) to calculate confidence intervals for all regression analyses. Bootstrapped CIs for NU and the three-way interaction did not contain zero. Furthermore, the adjusted *R*^2^ of the regression model was similar to the unadjusted *R*^2^, and the three-way interaction provided the same increase to adjusted and unadjusted *R^2^*, indicating that the regression model is likely unbiased by sample size or number of parameters (Montgomery & Morrison, 1973). Nonetheless, replication of the results demonstrated here in larger samples is essential. Another limitation of this study is the normative nature of our sample, as all participants were undergraduate students. Future work should examine whether these results replicate in clinically dependent, older, or community-based cohorts. However, college is an important time for development of problematic drinking behaviors (Rothman et al., 2008), and college students show high rates of harmful alcohol use (White & Hingson, 2014), so these results are important to the design of interventions targeting problematic drinking behaviors in college students.

### 4.5 Conclusion

We compared a model predicting clinically significant alcohol misuse by NU alone with a model predicting alcohol misuse with a combination of NU and two brain measures representing fundamental neurobiological mechanisms in addiction. We demonstrate that significant increases in predictive power of alcohol misuse are gained by the inclusion of neural correlates of cognitive control and reinforcement processing (the N2 and FRN ERP components) as moderating factors in the NU-alcohol misuse relationship. This suggests that interventions targeting cognitive control and reinforcement processing might reduce the risk impact of trait-levels variables on hazardous alcohol use.

## 4.7 Acknowledgments

Data were collected as part of ER’s dissertation at the University of Arkansas. Thanks to all volunteers who assisted in data collection, and particularly to those who supervised data collection sessions (in alphabetical order: Carroll Bentley, Zachary Levy, Stephanie Long, Morgan Middlebrooks, and Danielle Smith).

## 4.8 Data Availability

EEG data and stimulus presentation scripts described in this study are archived and publicly available at (https://osf.io/wu37k/).

## SUPPLEMENTAL MATERIAL for

In this supplement, we report more detailed control analyses for the moderated-moderation analysis described in the associated manuscript. Specifically, we present here the results of an analysis of positive reinforcement ERP amplitudes as moderators of the NU-AUDIT relationship, to assess whether our results are specific to negative reinforcement as predicted by theories of NU. The main results describe the use of raw condition ERP amplitudes as moderators in the NU-AUDIT relationship, as used in previous work from our group. However, as recent evidence has indicated that residualized ERP amplitudes might be applied as individual difference variables, we present here an analysis of the moderated-moderation analysis using ERPs that have been residualized by the opposing condition (i.e. incongruent N2 with congruent N2 regressed out, negative reinforcement better-than-expected FRN with worse-than-expected FRN regressed out).

The results of the positive reinforcement control analysis, summarized in Supplementary Table 1, demonstrated that in positive reinforcement conditions, N2 (cognitive contorl) but not FRN (reinforcement processing) moderated the NU-AUDIT relationship. This is as expected, given the theoretical importance of negative reinforcement processes in NU, and demonstrates that our results are specific to negative reinforcement outcome processing as opposed to positive reinforcement.

The results of the residualized ERP amplitude analysis, summarized in Supplementary Table 2, demonstrate that all of our key results, including most importantly the three-way interaction between NU, N2 amplitude, and FRN amplitude, were also significant when residualized ERP amplitudes were used rather than raw condition scores. This indicates that our analysis is not influenced by the correlated brain responses to congruent and incongruent trials, or by the correlated brain responses to better- and worse-than-expected negative reinforcement outcomes.

**Supplementary Table 1.** Given the theoretical link between NU and negative reinforcement processes, it was important to test whether our results were specific to the processing of negative reinforcement outcomes. We ran a moderated-moderation analysis as described in the associated manuscript but using positive reinforcement ERP amplitudes as opposed to negative reinforcement. This analysis demonstrated that in positive reinforcement conditions, N2 (but not FRN) amplitude moderated the NU-AUDIT relationship, demonstrating that the moderated-moderation result we describe in the main manuscript is specific to negative reinforcement processing as hypothesized.

| | Unstandardized $\beta$ | 95% CI | $t(55)$ | $p$-value |
| --- | --- | --- | --- | --- |
| **Negative Urgency (NU)** | 3.51 | [1.08, 3.99] | 3.51 | 0.001** |
| **Response Inhibition (N2)** | 0.07 | [-0.46, 0.60] | 0.26 | 0.79 |
| **Salience Attribution (FRN)** | -0.51 | [-2.33, 1.30] | -0.57 | 0.57 |
| **NU × N2** | -0.52 | [-0.93, -0.11] | -2.58 | 0.01* |
| **NU × FRN** | -0.17 | [-2.68, 2.33] | -0.14 | 0.89 |
| **N2 × FRN** | 0.12 | [-0.24, 0.48] | 0.68 | 0.5 |
| **NU × N2 × FRN** | -0.68 | [-1.60, 0.24] | -1.49 | 0.14 |

**Supplementary Table 2.** Given a recent report that residualized ERP amplitudes might be informative as individual difference measures (Meyer et al., 2017), we ran a control analysis using incongruent N2 with congruent N2 regressed out and better-than-expected FRN with worse-than-expected FRN regressed out. The pattern of results, and most critically the three-way interaction that out results in the main manuscript rest on, was similar to the regression analysis using raw condition ERP amplitudes.

| | Unstandardized $\beta$ | 95% CI | $t(55)$ | $p$-value |
| --- | --- | --- | --- | --- |
| **Negative Urgency (NU)** | 2.13 | [0.81, 3.45] | 3.24 | 0.002** |
| **Response Inhibition (N2)** | 1.47 | [-1.24, 4.18] | 1.09 | 0.28 |
| **Salience Attribution (FRN)** | -2.58 | [-5.24, 0.08] | -1.95 | 0.057 |
| **NU × N2** | -0.44 | [-3.43, 2.54] | -0.3 | 0.77 |
| **NU × FRN** | -2.99 | [-5.92, -0.07] | -2.06 | 0.045* |
| **N2 × FRN** | -5.93 | [-10.51, -1.35] | -2.6 | 0.01* |
| **NU × N2 × FRN** | -6.06 | [-12.06, -0.06] | -2.03 | 0.048* |

